# Pre-analytical processing of plasma and serum samples for combined proteome and metabolome analysis

**DOI:** 10.1101/2022.04.26.489520

**Authors:** Hagen M. Gegner, Thomas Naake, Aurélien Dugourd, Torsten Müller, Felix Czernilofsky, Georg Kliewer, Evelyn Jäger, Barbara Helm, Nina Kunze-Rohrbach, Ursula Klingmüller, Carsten Hopf, Carsten Müller-Tidow, Sascha Dietrich, Julio Saez-Rodriguez, Wolfgang Huber, Rüdiger Hell, Gernot Poschet, Jeroen Krijgsveld

## Abstract

Metabolomic and proteomic analyses of human plasma and serum samples harbour the power to advance our understanding of disease biology. Pre-analytical factors may contribute to variability and bias in the detection of analytes, especially when multiple labs are involved, caused by sample handling, processing time, and differing operating procedures. To better understand the impact of pre-analytical factors that are relevant to implement a unified proteomic and metabolomic approach in a clinical setting, we assessed the influence of temperature, sitting times, and centrifugation speed on the plasma and serum metabolomes and proteomes from six healthy volunteers.

We used targeted metabolic profiling (497 metabolites) and data-independent acquisition (DIA) proteomics (572 proteins) on the same samples generated with well-defined pre-analytical conditions to evaluate criteria for pre-analytical SOPs for plasma and serum samples. Time and temperature showed the strongest influence on the integrity of plasma and serum proteome and metabolome. While rapid handling and low temperatures (4°C) are imperative for metabolic profiling, the analysed proteome showed variability when exposed to temperatures of 4°C for more than 2 hours, highlighting the need for compromises in a combined analysis. We formalised a quality control scoring system to objectively rate sample stability and tested this score using external data sets from other pre-analytical studies.

Stringent and harmonised standard operating procedures (SOPs) are required for pre-analytical sample handling when combining proteomics and metabolomics of clinical samples to yield robust and interpretable data on a longitudinal scale and across different clinics. To ensure an adequate level of practicability in a clinical routine for metabolomics and proteomics studies we suggest to keep blood samples up to 2 hours on ice (4°C) prior to snap-freezing as a compromise between stability and operability. Finally, we provide the methodology as an open source R package allowing the systematic scoring of proteomics and metabolomics datasets to assess the stability of plasma and serum samples.

## Introduction

Mass spectrometry-based metabolomics and proteomics are emerging technologies that are increasingly employed in laboratory and clinical settings to refine our understanding of disease biology, vulnerabilities, and resistance mechanisms. Liquid biopsies, such as blood, provide the opportunity to collect information on a patient’s metabolome and proteome status on a longitudinal scale to track disease progression or response to a treatment (Tsonaka *et al*., 2020; Gummesson *et al*., 2021). For instance, longitudinal metabolomic profiling of plasma collected from patients suffering from COVID-19 was linked to disease progression, including a panel of metabolites collected at the onset of the disease that may predict the disease severity (Sindelar *et al*., 2021). Similarly, proteomic analysis of COVID-19 patients revealed protein signatures associated with survival, tissue-specific inflammation, and disease severity (Filbin *et al*., 2021). The independent analysis of such complex diseases yields promising findings, highlighting that the present technologies are not the limiting factors for the broader use of mass spectrometry (MS) in clinical workflows.

MS-based technologies have matured over the past years, allowing the investigation of analytically challenging but highly informative samples such as blood plasma and serum. Technical advances comprise of but are not limited to: i) Increased reproducibility and automation in sample preparation, ii) faster, more sensitive, and robust MS instruments, and iii) improved data analysis algorithms, multi-omics and integrative workflows. While these developments reduce technical noise in the data sets and improve the detection of true biological variability, their efficacy may be compromised if the quality of the starting material is not strictly controlled and standardised.

Although standard operating procedures (SOPs) for blood collection are often in place to suit clinical routine, they may not be harmonised between clinics, and they usually are not optimised to preserve proteins and metabolites for subsequent *omics* analyses. In particular, differences in sample handling (e.g., temperature, sitting time, use of anticoagulants) may alter the observable protein and metabolite patterns. In biomarker discovery studies, these pre-analytical factors are crucial and have to be considered by clinicians and analysts (Lippi *et al*., 2020).

Previous studies have highlighted the effects of such pre-analytical factors and recommend best practices often for metabolomic analyses (Yin, Lehmann and Xu, 2015) or proteomic analyses (Hassis *et al*., 2015) independently. While either technique already produced a suite of potential quality markers related to blood samples, to our knowledge, few studies analysed the effect sizes of varying pre-analytical parameters in a combined proteomic and metabolomic analysis on the same samples, to harmonise the requirements for both techniques with such a comprehensive set of features (Cao *et al*., 2019). Critically, sample collection and handling requirements differ between metabolomics and proteomics and need to be adjusted accordingly for a combined clinical SOP.

Here, we assess how pre-analytical factors impact on metabolite and protein levels in plasma and serum samples caused by differences in sitting time, temperature regimes (4°C room temperature (RT), only RT for serum), and centrifugal acceleration levels. Using targeted metabolic profiling and a single-shot, data-independent acquisition (DIA) proteomics approach, we determine that keeping blood samples on ice (4°C) for up to 2 hours prior to snap-freezing are the optimal conditions to preserve metabolites and proteins for a combined metabolomics/proteomics workflow. We introduce an open-source scoring system to assess the quality of plasma and serum samples (Figure 1B).

## Results

To assess how sample handling and treatment affects the stability of protein and metabolite levels in human plasma and serum samples, we selected four time points between centrifugation and snap-freezing to quench samples, as follows: 0 h as the baseline of metabolite and protein levels immediately after sampling, 2 h as the clinically feasible time point to quench samples, 4 h as a middle point, and 8 h (quenching at the end of a typical working day) (Figure 1A). Furthermore, samples were kept at different temperatures during these sitting times (on ice/4°C and at RT) to investigate their influence on altering metabolome and proteome composition. Additionally, we included two centrifugation schemes for plasma samples (2000 g and 4000 g). However, we could not attribute any significant changes in the plasma metabolome and proteome between different centrifugation conditions, and therefore only applied 2000 g in the following sections.

**Figure 1:**
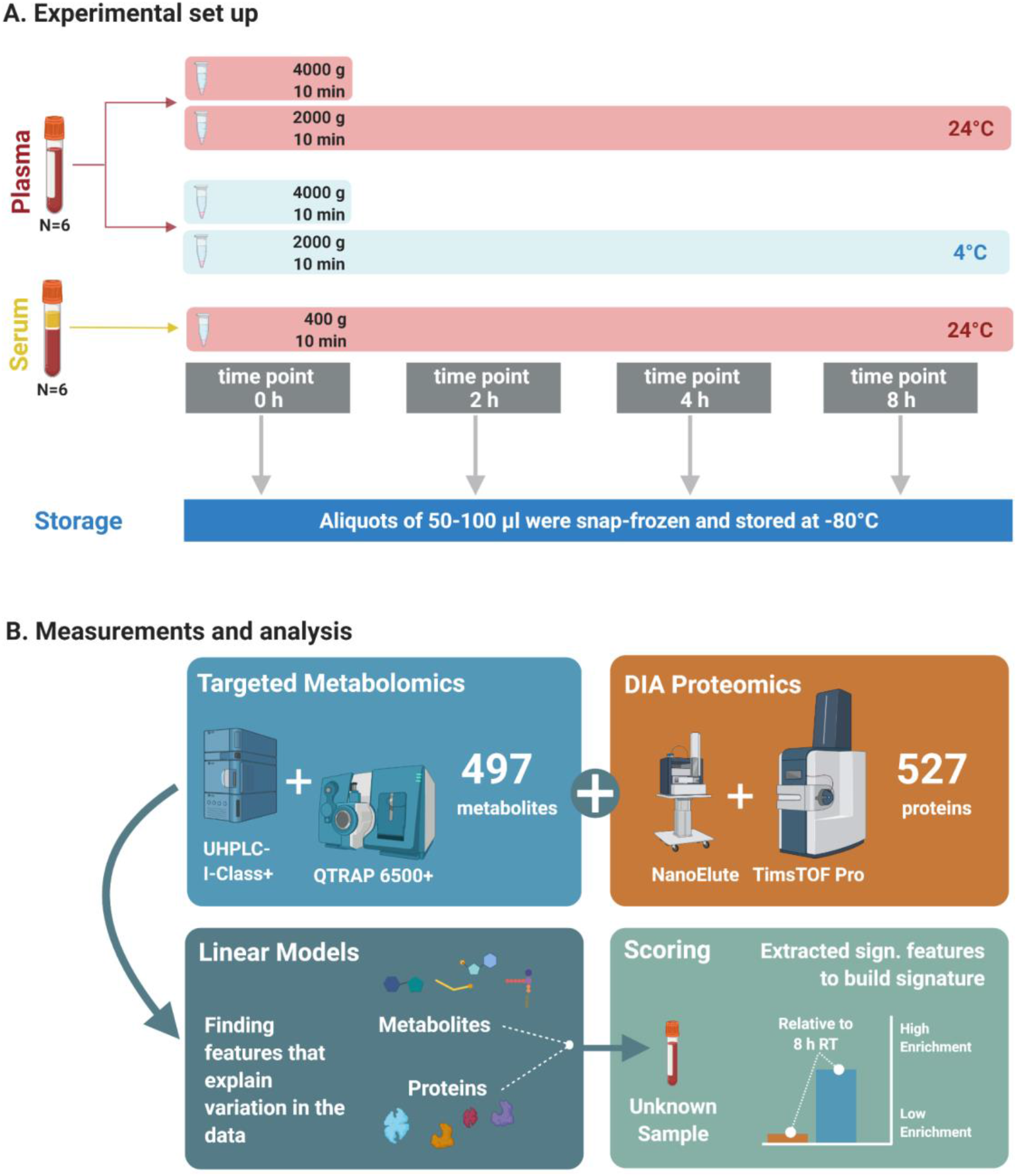
Experimental setup and analyses to assess effects of sample handling or treatment on metabolite and protein levels. A: Plasma and serum samples from six healthy individuals were subjected to different sample treatment and handling conditions: sitting time of 0, 2, 4, and 8 h; incubation temperature of 4°C and 24°C (RT); and two different centrifugal levels (2000 and 4000 g) for the plasma samples. Due to quality reasons, the metabolomics data set only consisted of data from five individuals (see Methods). B: Following the factorial outline, metabolite and protein levels were obtained by mass spectrometry. Next, proteomic and metabolomic data were analysed using linear models. The significant features were extracted and used to build scores to assess sample quality.

### Identifying metabolites and proteins affected by temperature and sitting time

Analysis of plasma and serum samples by targeted metabolic profiling and a single-shot, data-independent acquisition (DIA) proteomics yielded quantitative information for in total 497 metabolites and 572 proteins. An initial LIMMA analysis (Table 1) showed a high number of features that differed between individual blood donors (α < 0.05, after FDR correction using the Benjamini-Hochberg, BH, method). In addition, PCA (Supplementary Figure 1A) of the proteomic and metabolomics data sets indicated individual-specific effects, a finding that was further supported by t-SNE and UMAP analyses (Supplementary Figure 2) and multi-omics factor analysis on both data modalities (MOFA, Supplementary Figure 3). Especially for the metabolomics data set there was a clear separation between individuals, while for the proteomics data set we found less pronounced effects (Supplementary Figure 2D). This analysis suggests that there are dominating individual effects that are reflected in the metabolomics data set and to a lesser extent in the proteome. To gain further insight into this, we next performed classification by Partial least square – discriminant analysis (PLS-DA) and sparse PLS-DA (sPLS-DA) to discriminate the individuals based on the metabolite and proteomics profiles. Indeed, it was possible to classify the individuals using the metabolite and protein levels with a low classification error (Supplementary Figure 1C), suggesting that there are features in the metabolomics and proteomics data where individual-specific effects are prevalent. The metabolites and proteins in Supplementary Table 1 were selected by sPLS-DA to explain the variance using the *individual* as the class vector (Supplementary Figure 1B).

**Table 1:** Number of features with a significant effect from the factor individual. Shown is the number of significant features after FDR (α < 0.05). The total number of features are 497 (metabolites) and 572 (proteins).

|  | plasma | serum |
| --- | --- | --- |
| metabolites | 470 | 435 |
| proteins | 221 | 151 |

We also performed PLS-DA to discriminate for *time* and the combination of *time* and *temperature* (Supplementary Figure 4). Both binary classification problems yielded models with high classification rate error and lower values for the explained variance for both the proteomics and metabolomics data set (Supplementary Figure 4A and C) compared to *individual* as the class vector (Supplementary Figure 1). The sPLS-DA analysis yielded a list of features that were used to explain the class vectors of the data and could be regarded as features that change along the *time* and *time*/*temperature* axes (Supplementary Figure 4B and D). We included this list as Supplementary Tables 2, 3, and 4.

In a next step, we looked into the changes of metabolite and protein levels when considering inter-individual differences. Motivated by the results of the previous analyses (initial LIMMA analysis, dimension reduction analysis, PLS-DA, MOFA), showing that metabolome and proteome variation is influenced by inter-individual differences, we decided to use mixed linear models to determine the features that will change according to sitting time, temperature or a combination of time and temperature. We modelled as fixed effects *time*, *temperature*, and the interaction term *time/temperature* (plasma) and *time* (serum). The information on the blood donor (*individual)* was included for both groups as a random effect (Figure 2A). An overview of the absolute change of the significant metabolites and proteins can be found in Figure 2B (see also Figure 2C for exemplary metabolites and proteins). We provide the metabolite- and protein-associated p-values for plasma and serum samples in the Supplementary Information.

**Figure 2:**
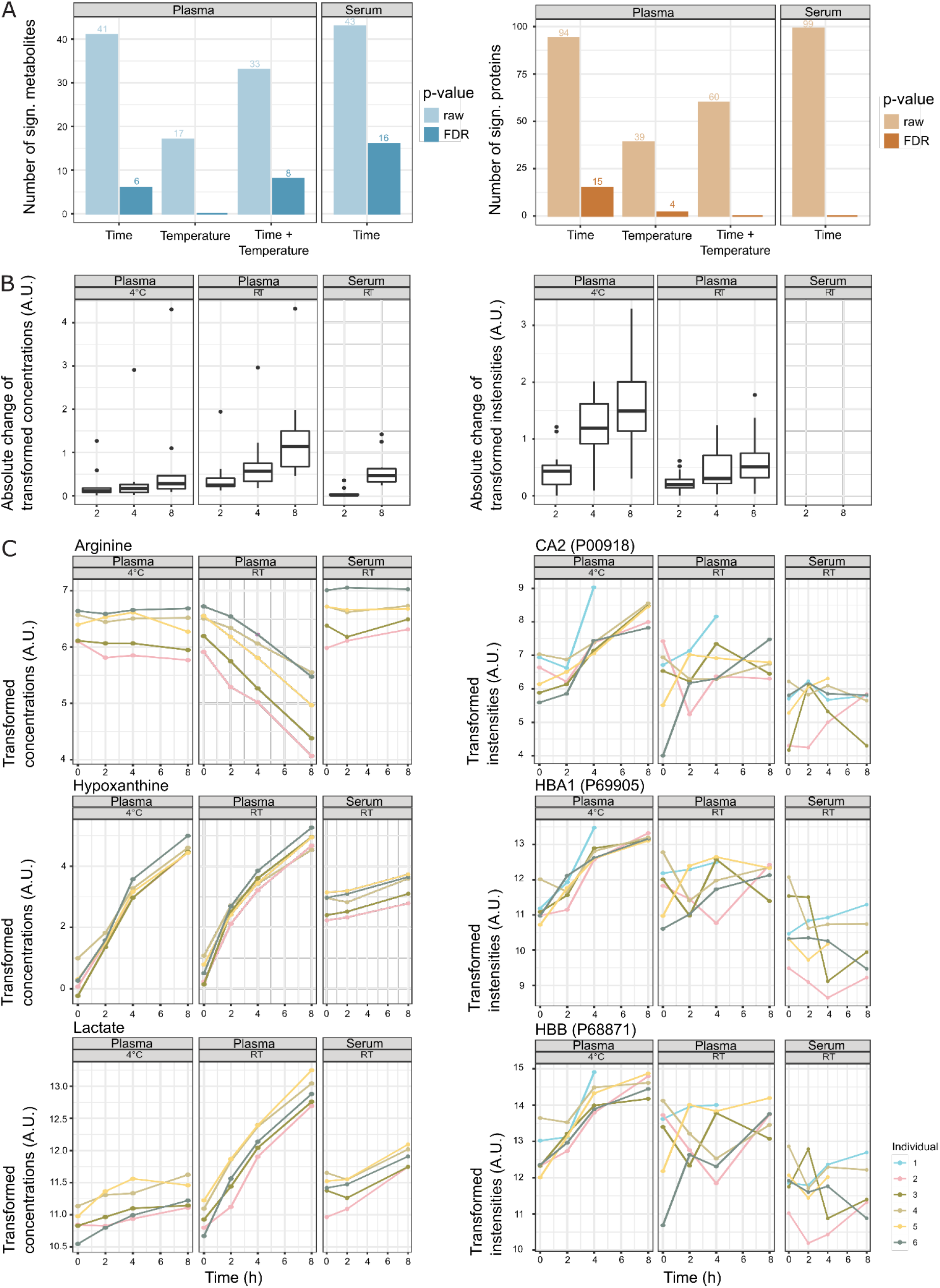
Stability of metabolites and proteins along the sitting time and temperature axes. A: Number of significant metabolites or proteins that show changes according to pre-analytical factors *time*, *temperature* and the interaction *time/temperature* (raw p-values and p-values after FDR correction, α < 0.05). B: Absolute change of transformed concentrations/intensities of significant metabolites and proteins along the time axis (after FDR correction). The changes to the time point 2 h, 4 h, and 8 h are displayed as the mean changes of the individuals (intensity at time point 0 h is 0). For plasma, the features are included that are significant for the pre-analytical factors *time*, *temperature* or the interaction *time/temperature* (α < 0.05, FDR correction). For serum, the features are included that are significant to the pre-analytical factor *time* (α < 0.05, FDR correction). C: Examples of metabolites and proteins that show a significant association with the pre-analytical factor *time* (hypoxanthine, lactate, CA2, HBB, HBA1) or interaction *time/temperature* (lactate, arginine) (α < 0.05, after FDR correction).

Looking at metabolomics- and proteomics-specific differences, the analysis revealed that metabolite concentrations were less stable at RT, while protein abundances were less stable at 4°C (Figure 2 and Supplementary Figure 5). For the affected features the absolute change was in most cases more prominent after 8 h than 2 h, yet they were not significant (Supplementary Figure 5).

### Scoring plasma and serum sample quality using proteomic and metabolomic signatures

We next investigated whether patterns of potential protein and metabolite deregulation (with respect to *time* and *temperature*) could be used as a quality metric for samples obtained under the tested conditions. We selected the top 20 proteins and metabolites ranked by p-value to generate signatures of the following handling conditions: plasma kept on ice (4°C) or RT for 8 hours, and serum at RT for 8 hours (Supplementary Figure 6). While it may be difficult to draw conclusions regarding the significance of individual metabolites or proteins due to limited sample size, their combined signal may hold enough information to score the relative quality of samples with respect to sitting time and temperature. Thus, to confirm that these signatures could yield sensible insight into sample integrity, we computed an average normalised enrichment score (NES, see Methods) for each signature in the respective sample pre-analytical conditions. If the signatures are indeed informative, we should expect to observe a steady increase of enrichment of the respective signature for each condition. Coherently, each signature showed higher scores for samples that matched the respective conditions from which they were derived (Figure 3A). This showed notably that the plasma protein signature at RT over time already scored highly in samples stored at RT for 4 h, as opposed to the signature of plasma/4°C/8 h which only scored high in the samples obtained at low temperature and after 8 h, as expected. This pattern was inverted for metabolic signatures of plasma samples. This indicates that the changes are more pronounced at the metabolomic level when samples are stored at 4°C compared to RT, while changes are more pronounced at the proteomic level for plasma samples kept at RT.

**Figure 3:**
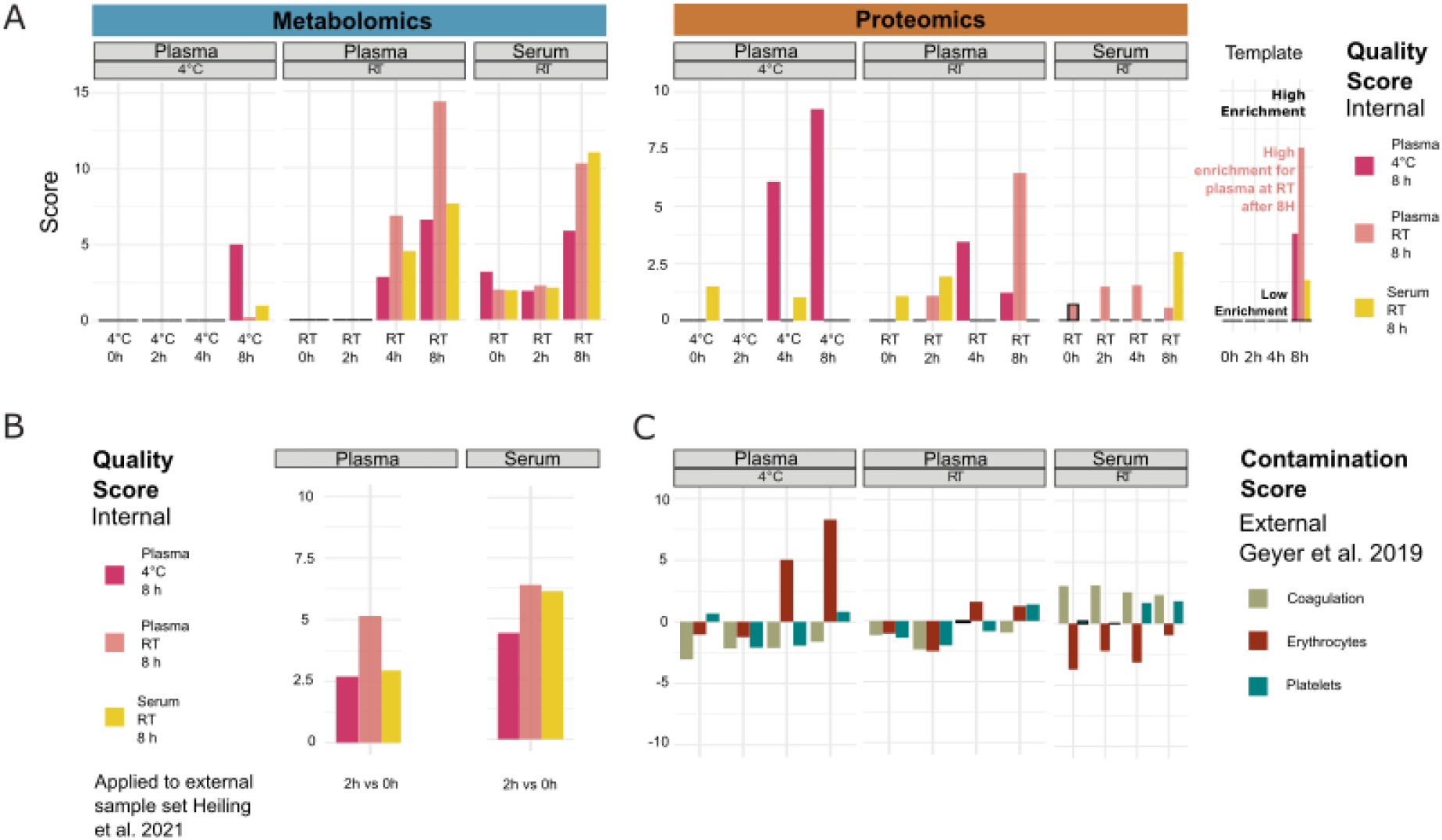
Systematic evaluation of sample quality and contamination with proteomic and metabolomic signatures. A: Proteomic/Metabolic pre-analytical condition signature scores for each sample group (as averaged from individual samples). Signature scores are normalised enrichment scores, representing the number of standard deviations away from the mean of an empirical null distribution of scores. A high score means that the sample displays an enrichment of markers of the corresponding signature, compared to other samples. While the scores generated can be both positive and negative, we focus exclusively on positive scores in this figure. B: Metabolic pre-analytical signature applied to score samples from an external study (Heiling *et al*., 2021). C: Coagulation, erythrocyte, and platelet signatures were used to score contamination of proteomic plasma and serum samples. Signatures were taken from (Geyer *et al*., 2019).

Finally, while the NES can take both positive and negative values, here, we focused only on the positive values to simplify the interpretation of the results. Since the data was normalised in a way that each measurement is scaled relative to other samples of the cohort, the scores will be drawn from a distribution where a NES of 0 represents samples that have average levels of degradation compared to the overall cohort, and any value above that represent samples that show higher degradation compared to the rest of the cohort. It is worth noting that this scoring can only score samples in a relative manner to the rest of the cohort, and cannot provide absolute quantification of sample degradation.

To validate this approach, we used the signatures to score metabolomic results from an external study (Heiling *et al*., 2021), where plasma and serum samples were kept at RT for 2 h (Figure 3B). The plasma/RT/8 h metabolic signatures got a higher score (NES = 5.1) than plasma/4°C/8 h (NES = 2.6) and serum/RT/8 h (NES = 2.9) signatures. However, the serum/RT/8 h signature score was similar to the plasma/RT/8 h signature score (NES = 6.1 and NES = 6.4, respectively). Thus, the serum RT/8 h and plasma/RT/8 h metabolic signature appears less discriminant than the plasma/4°C/8 h metabolic signature. Nevertheless, the best scores overall matched with the actual experimental conditions that were used, indicating that the scoring system holds beyond the data set that was used for training.

In order to further characterise the changes that we observed in the plasma and serum samples over time, we investigated if proteomic signatures could be associated with contamination by proteins originating from specific blood cells. We obtained proteomic markers of coagulation, erythrocyte, and platelet contamination from (Geyer *et al*., 2019) Plasma samples kept on ice (4°C) for 4 h and 8 h showed the highest enrichment of erythrocyte contamination markers (Figure 3C) mainly driven by CAT, CA2, BLVRB, PRDX2, and ALDOA (Supplementary Figure 6A, B). Interestingly, the plasma/4°C/8h seems to be also specifically driven by a lower abundance of the VWF protein, a blood glycoprotein involved in platelet adhesion (Supplementary Figure 6C). The contamination scores were lower in plasma samples that were kept at RT, although they still showed a progressive increase over time. On the other hand, serum samples exhibited no significant increase in erythrocyte contamination score, instead showing a consistently high (albeit slightly decreasing) score for coagulation markers over time, as expected. This signature was mainly driven by increased PPBP and THBS1 and decreased F13A1 (Supplementary Figure 6A and D). In a similar fashion, we displayed the main drivers of the metabolomic-derived signatures such as hypoxanthine, lactate, ornithine and aspartate (Supplementary Figure 6A, E, F, G). Taken together, these results show that the scoring provides a quantitative metric for the quality control for proteomic and metabolomic data of plasma and serum samples. This should be a helpful tool to exclude low-scoring samples for further analysis.

## Discussion

The progress of MS-based technologies over the past years has enabled the characterization and quantification of analytically challenging but clinically accessible samples such as blood plasma and serum. Although SOPs for individual metabolic and proteomic analyses have been developed (Pasella *et al*., 2013; Yin, Lehmann and Xu, 2015; Tuck, Turgeon and Brenner, 2019; Lippi *et al*., 2020), there is no consensus on their combined application for the molecular characterization of blood samples in a clinical setting.

Here, we performed a comprehensive analysis on the stability of 497 and 572 metabolites and proteins in blood plasma and serum to scrutinise the effects of various treatment regimes (sitting time and incubation at different temperatures) to simulate different sample handling scenarios. Notably, our experiment aimed to define an SOP trade-off regarding the different requirements for metabolomics and proteomics, to effectively apply both approaches to the same blood samples. In addition, we aimed to implement an objective quality scoring as a metric for sample quality and potential contamination. Although this study was performed on a small cohort of healthy volunteers, the findings have implications on the sampling procedure of clinical blood collection as well as the bioinformatics analysis for quality control.

### Measuring the change of the metabolome and proteome

By our statistical analyses, we detected changes in numerous features that may affect the biological interpretation of clinical metabolomics and proteomics data sets. Some metabolites (e.g. hypoxanthine, xanthine, lactate, arginine, ornithine, cystine) and proteins (e.g. CA1, CA2, HBB, HBD, HBA1) showed a profound dependency on sitting time and temperature (or a combination thereof) (Figure 2 and Figure 4A).

**Figure 4:**
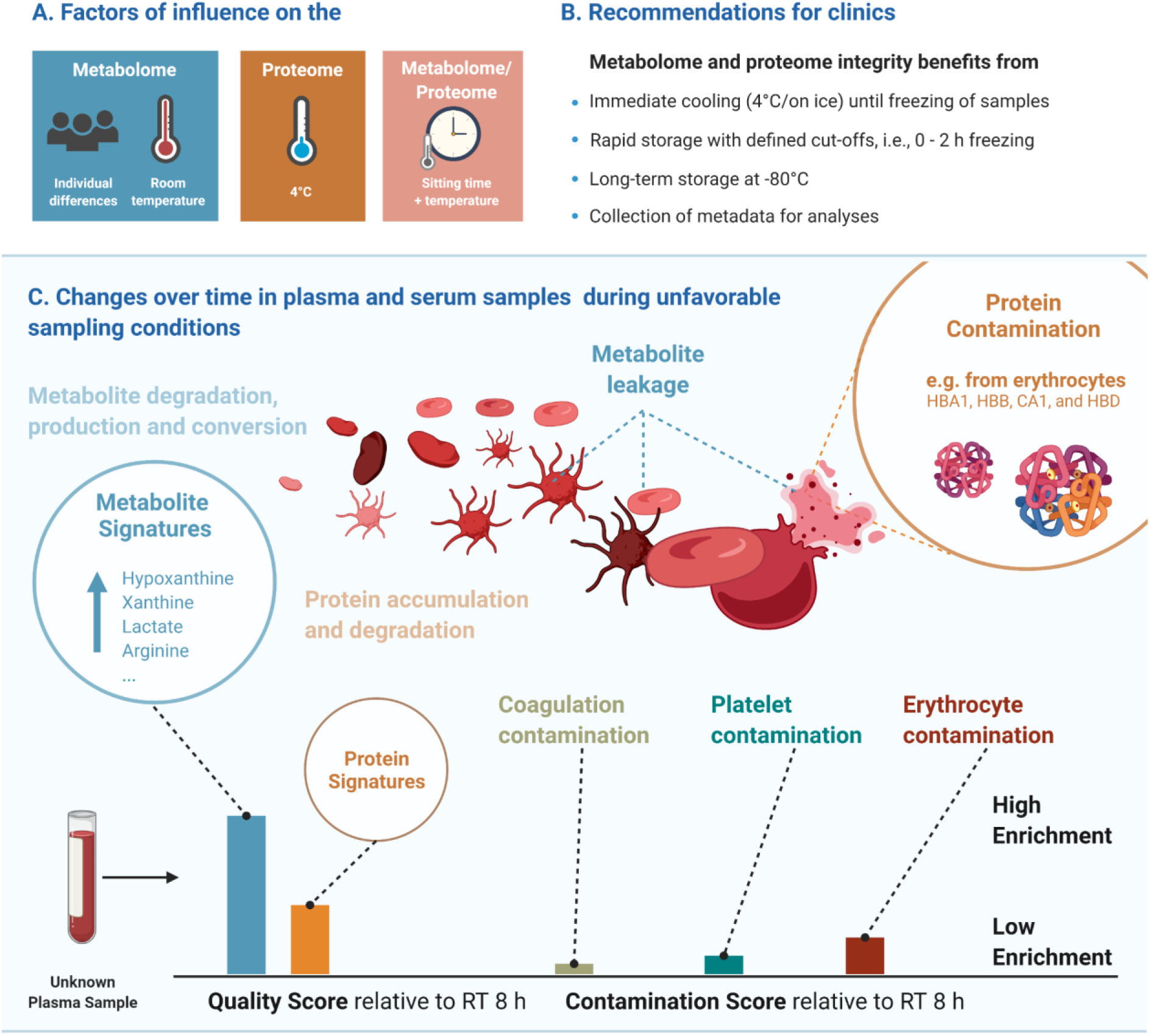
Considerations for the joint proteomic and metabolomic analysis of plasma and serum samples. A: Most influential factors on the proteome and proteome in our data. B: Recommendations for clinical blood sampling. C Change over time in blood plasma and serum samples and the resulting quality and contamination scores.

Metabolite classes such as amino acids, purines, and carbohydrates vary in abundance and are therefore usually investigated to answer biological questions. Yet, these well-known metabolites affected by the conditions represent only 10% of our data set, and we have increased the panel of temperature-time sensitive metabolites due to the broad metabolite coverage. In fact, lipids are the largest observed class of metabolites, most of which were stable across the tested conditions (Supplementary File 1).

Temperature and time are well known to affect metabolite and protein levels (Kamlage *et al*., 2014; Cao *et al*., 2019; Daniels *et al*., 2019; Stevens *et al*., 2019). Elevated levels of hypoxanthine and amino acids over time (Ferreira *et al*., 2019) and the deregulation of cholesterol metabolism were also previously documented (Ryu *et al*., 2016). Association of these and other metabolites to a pathological condition, therefore, needs to be evaluated with caution, to exclude that they emerge inadvertently by sample handling or technical bias. Strict pre-analytical measures help to gain confidence in the subsequent biological and clinical interpretation based on the measured features.

For most of the features in our data sets, we only found minor changes under the experimental conditions applied (Figure 2). There, 90% of metabolites and 97% of proteins only varied slightly over time. Other metabolome studies reported similar proportions where 91% of the metabolite remained stable over several pre-analytical conditions (Ferreira *et al*., 2019). This implies that in clinical research studies where large effect sizes from biological differences are known or expected and large cohorts were used, the contribution to feature level variation stemming from sample handling might be partially alleviated. The integration of several data sets, i.e., proteomic and metabolomic, from the same sample may also mitigate bias.

### Rapid handling and cold storage of up to 2 h as SOP

Although the increased stability of the metabolome at 4°C was expected, we observed a contrary effect for proteins, showing higher variance at low temperature (Figure 2 and 3A), suggesting that proteomics and metabolomics require different pre-analytical conditions to obtain optimal results. Therefore, we propose that keeping plasma and serum samples on ice for up to 2 h is an acceptable trade-off to maintain adequate stability of both the proteome and metabolome (Figure 2 and Supplementary Figure 5). In addition, this should be a condition that can be conformed to in clinical practice (Figure 4B).

Similarly, considerations may be made regarding the storage time of samples in biological repositories such as biobanks. Previous studies showed that long-term storage at -80°C over seven years only introduces minimal variation, and that significant changes occur upon longer storage times (Wagner-Golbs *et al*., 2019). This highlights the potential to address clinical questions using metabolomics and proteomics from biological repositories under the prerequisite that the sampling collection is comparable. While biobank samples are an essential resource for discovery studies, prospective samples enable the enforcement of SOPs during collection that are more suitable for metabolomic analyses, e.g., by storing samples at 4°C for under 2 hours and then quench by snap-freezing in liquid nitrogen.

### Quality control signatures to score plasma and serum samples

Designing formal criteria for data curation and analysis is crucial to ensure data robustness. To this end, we devised a scoring system using the significantly altered proteins and metabolites as signatures to evaluate the impact of pre-analytical conditions on proteome and metabolome integrity of a given sample. Provided as an R package (https://github.com/saezlab/plasmaContamination), this tool can be used for quality control after pre-analytical handling, and in addition, the proteome signatures enable to distinguish the severity and the source of contamination, i.e. from platelets, erythrocytes or resulting from coagulation (Figure 3C and 4C). We showed that the changes in protein abundance in samples stored on ice were mainly related to protein markers of erythrocytes in plasma samples, likely resulting from hemolysis occurring under this condition (Figure 4C). As expected, coagulation signatures scored exclusively high in serum samples.

Both the scores for metabolite and protein contamination enable the quality assessment of plasma and serum samples of unknown origin. Of note, the erythrocyte, platelet, and coagulation signatures were obtained from a large external cohort of samples (>70 samples), while the signatures derived from our own samples were estimated from a comparatively small number of samples (n=6). Although this may affect their discriminative power, the derived signatures and bioinformatic tools are publicly accessible and can therefore be updated and expanded easily when more data become available. Still, those signatures yielded coherent scores when they were tested with our own samples and were validated with samples from an external study. The expansion of such signatures towards other pre-analytical factors, such as storage conditions, enables the development of further quality control metrics. This may be achieved through similarly structured experimental set-ups with small sample sizes or the analysis of bigger cohorts with the inclusion of metadata. We anticipate that a quality score for proteome and metabolome integrity can have great practical utility, enabling the exclusion of low-scoring samples for further analysis. This will be particularly important if clinical decisions are to be made based on metabolic or proteomic data from such samples. At this point, it is premature to suggest a cut-off score here, since the number of samples in our study is low, and since the choice for such a cut-off may depend on the setting of the analysis (e.g., biomarker discovery, clinical decision). Finally, although a quality score is helpful, it cannot replace rigorous SOPs. In addition, this must be evaluated in the context of other available metadata that should be applied in combination with other quality control strategies (Naake and Huber, 2022).

### Summary

In this study we assessed the influence of controllable pre-analytical parameters on protein and metabolite levels in plasma and serum samples, to define or improve SOPs for concerted metabolomic and proteomic analyses. While only a subset of metabolites and proteins changed, the ability to identify features that are prone to alteration increases the confidence in such broadly acquired data sets. We propose to store blood samples for maximum 2 h on ice (4°C) before quenching the samples, as a compromise between stability and practical operability. Additionally, the metabolomic and proteomic signatures can be applied routinely in bioinformatics workflows to review and evaluate the sample quality of plasma and serum samples. Due to its accessibility, such signatures may be expanded over time to improve the assessment of qualitative differences between blood samples. Lastly, bigger sample sizes and additional metadata of volunteers and/or available metadata from clinics may extend these scores to include signatures capturing other sources of variability important to clinical studies, such as storage, medication, or lifestyle.

## Methods

### Sampling and sample treatment /design

Peripheral blood samples were collected from 6 male healthy volunteers (age 22-37 years, median age 29.83 years) with written informed consent in accordance with the Declaration of Helsinki and approved by the Ethics Committee of the Medical Faculty of the University of Heidelberg (S-254/2016). All samples were taken in the early morning in fasting condition using Serum and EDTA S-Monovette tubes (Sarstedt AG, Nümbrecht, Germany). Serum samples were allowed to coagulate for 30 min after collection, were kept at RT for the indicated time points and centrifuged at 400 x g for 10 min. Plasma samples were kept at RT or at 4°C/on ice for the indicated time points, followed by centrifugation for 10 min. at 2000 x g and 4000 x g, respectively. Subsequently, samples were divided into single-use aliquots, snap-frozen in liquid nitrogen and stored at -80°C until analysis. Thus, metabolomic and proteomic analyses were performed from the same original samples.

### Metabolomics

For the metabolomics analysis of up to 630 metabolites the Biocrates MxP® Quant 500 kit (Biocrates, Austria) was used following the manufacturer’s protocol. Briefly, 10 µl of plasma or serum was semi-automated pipetted on a 96 well-plate containing internal standards using a pipetting robot (epMotion 5057, Eppendorf, Germany) and subsequently dried under a nitrogen stream using a positive pressure manifold (Waters, Germany). Afterwards, 50 µl phenyl isothiocyanate 5% (PITC) was added to each well to derivatize amino acids and biogenic amines. After 1 h incubation time at RT, the plate was dried again. To resolve all extracted metabolites 300 µl 5 mM ammonium acetate in methanol were pipetted to each filter and incubated for 30 min. The extract was eluted into a new 96-well plate using positive pressure. For the LC-MS/MS analyses 150 µl of the extract was diluted with an equal volume of water. Similarly, for the FIA-MS/MS analyses 10 µl extract was diluted with 490 µl of FIA solvent (provided by Biocrates). After dilution, LC-MS/MS and FIA-MS/MS measurements were performed in positive and negative mode on subsequent days. For chromatographic separation an UPLC I-class PLUS (Waters, Germany) system was used coupled to a QTRAP 6500+ mass spectrometry system (Sciex, Germany) in electrospray ionisation (ESI) mode. Data was recorded using the Analyst software suite (version 1.7.2, Sciex, Germany) and transferred to the Met*IDQ* software (version Oxygen-DB110-3005, Biocrates, Austria) which was used for further data processing, i.e., technical validation, quantification and data export.

Low-abundant metabolites that were not measured in more than 66% of the samples as 10-times the levels over the limit of detection (LOD) or above the lower limit of quantification (LLOQ) (both according to the MetIDQ software) were removed from the subsequent analysis. Using the MatrixQCvis package (Naake and Huber, 2021; version 1.1.0), low-quality samples were removed leading to the exclusion of all samples belonging to individual 1 from the further analysis. To prepare the data sets for statistical analysis, the raw intensity values were transformed via vsn (vsn2 function from vsn package, version 3.59.1). which yields a matrix with feature values being approximately homoskedastic (features have constant variance along the range of mean values). Missing values were imputed via the impute.MinDet function (imputeLCMD package, version 2.0). While for MOFA analysis, the transformed data set was used, for all other analyses (dimension reduction, PLS-DA, limma analyses, mixed linear models), the imputed data set was used.

### Proteomics

#### Sample preparation

Plasma and serum aliquots were diluted 1:10 in ddH2O to perform a bicinchoninic acid assay (BCA, Pierce – Thermo Scientific) protein quantification. Subsequently, 10 μg protein per sample were further processed in a 1 μg/μL concentration and 100 mM ammonium bicarbonate (ABC, Sigma-Aldrich) using a Covaris LE220plus for AFA-based ultrasonication in a 96-well format. The plate was transferred to a Bravo pipetting system (Agilent Technologies, USA) for autoSP3 processing as described elsewhere (Müller *et al*., 2020). In brief, 10 mM TCEP, 40 mM chloroacetamide (CAA), 100 mM ABC, and 1x protease inhibitor cocktail (PIC, cOmplete, Sigma-Aldrich) were added to each sample, followed by incubation at 95°C for 5 minutes. Protein binding to Sera-Mag Speed Beads (Fisher Scientific, Germany) was induced by increasing the buffer composition to 50% acetonitrile (ACN, Pierce – Thermo Scientific). The bead stock was prepared as follows: 20 μL of Sera-Mag Speed Beads A and 20 μL of Sera-Mag Speed Beads B were combined and rinsed with 1x 160 μL ddH2O, 2x with 200 μL ddH2O, and re-suspended in 20 μL ddH2O for a final working stock. The bead stock was prepared for all samples. The autoSP3 protein clean-up was performed with 2x ethanol (EtOH, VWR International GmbH, Germany) and 2x ACN washes. Reduced and alkylated proteins were digested on-beads and overnight at 37°C in a lid-heated PCR cycler (CHB-T2-D ThermoQ, Hangzhou BIOER Technologies, China) in 100 mM ABC with sequencing-grade modified trypsin (Promega, USA). Upon overnight protein digestion each sample was acidified to a final concentration of 1% trifluoroacetic acid (TFA, Biosolve Chimie). MS injection-ready samples were stored at -20°C.

#### Data acquisition

Peptide samples were measured using a timsTOF Pro mass spectrometer (Bruker Daltonics, Germany) coupled with a nanoElute liquid chromatography system (Bruker Daltonics, Germany). Peptides were separated using an analytical column (Aurora Series Emitter Column with CSI fitting, C18, 1.6 μm, 75 μm x 25 cm) (Ion Optics, Australia). The outlet of the analytical column with a captive spray fitting was directly coupled to the mass spectrometer using a captive spray source. Solvent A was ddH2O (Biosolve Chimie), 0.1% (v/v) FA (Biosolve Chimie), 2% acetonitrile (ACN) (Pierce, Thermo Scientific), and solvent B was 100% ACN in ddH2O, 0.1% (v/v) FA. The samples were loaded at a constant maximum pressure of 900 bar. Peptides were eluted via the analytical column at a constant flow of 0.4 μL per minute at 50°C. During the elution, the percentage of solvent B was increased in a linear fashion from 2 to 17% in 22.5 minutes, then from 17 to 25% in 11.25 minutes, then from 25 to 37% in a further 3.75 minutes, and then to 80% in 3.75 minutes. Finally, the gradient was finished with 3.75 minutes at 80% solvent B. Peptides were introduced into the mass spectrometer via the standard Bruker captive spray source at default settings. The glass capillary was operated at 3500 V with 500 V end plate offset and 3 L/minute dry gas at 180°C. Data were acquired in data-independent acquisition (DIA) mode using full scan MS spectra with mass range m/z 100 to 1700 and a 1/k0 range from 0.6 to 1.6 V*s/cm^2^ with 100 ms ramp time were acquired with a rolling average switched on (10x). The duty cycle was locked at 100% and the TIMS mode was enabled. All timsTOF Pro and nanoElute methods were default provided by Bruker. Data were acquired in data-independent acquisition (DIA) mode using

#### DIA method details

For the DIA scans, resolution was set to 30,000 FWHM, with an automatic gain control (AGC) target of 3 x 10^6^ ions, a fixed first mass of 200 m/z, a stepped collision energy of 27, and a loop count of 34 with an isolation window of 24.3 m/z.

#### Data processing

Raw files were processed in Biognosys Spectronaut version 14.11. The search parameters were set to default as specified by the developer of the software. In brief, enzyme was set to trypsin/P with up to 2 missed cleavages. Carbamidomethylation (C) was selected as a fixed modification; oxidation (M), acetylation (protein N-term) were set as variable modification.

Data quality was checked by the MatrixQCvis package (Naake and Huber, 2021, version 1.1.0), leading to the exclusion of several low-quality samples. Further data processing was done according to the metabolomics data set using vsn transformation and imputation of missing values by the impute.MinDet function.

### LIMMA analysis to test for *individual* effects

For the metabolomics and proteomics data set, the transformed intensities were taken. Separately for the plasma and serum samples, a linear model was fitted to the data (using lmFit from limma, version 3.50.0). For plasma samples, information on the *individual*, *time*, *temperature* and the interaction between *time* and *temperature* were included as terms into the model. For serum samples, information on the *individual* and *time* were included as terms into the model. t-statistics and moderated F-statistics were computed by empirical Bayes moderation of the standard errors towards a global value (using eBayes from limma). The corresponding p-values to the effects for all individuals were adjusted via FDR using the Benjamini-Hochberg method (α < 0.05). The code for the analysis can be found here: https://github.com/tnaake/SMARTCARE_preanalytical_processing/tree/main/LIMMA

### Multi-omics factor analysis (MOFA) of the combined metabolomics and proteomics data sets

The vsn-transformed data sets (no imputed missing values) were used for running MOFA. For the proteomics data set, the mean intensities between duplicates were calculated. Subsequently, only the overlapping samples (intersection) of the metabolomics and proteomics data sets were retained (40 plasma samples and 14 serum samples). MOFA (using the MOFA2 package, version 1.1.21) was run using the metabolomics and proteomics data sets as views and Plasma and Serum as groups. The data options were set to default values (scale_views = FALSE, scale_groups = FALSE, center_groups = TRUE, use_float32 = FALSE), the model options were set to default (gaussian likelihood for views, maximum number of factors = 15, spikeslab_factors = FALSE, spikeslab_weights = TRUE, ard_factors = TRUE, ard_weights = TRUE), and the training options were set to default (maximum of iterations = 10000, convergence mode = „slow“, drop_factor_threshold = 0.01, startELBO = 1, freqELBO = 5, stochastic = FALSE, gpu_mode = FALSE, seed = 42, weight_views = FALSE). The code for the analysis can be found here: https://github.com/tnaake/SMARTCARE_preanalytical_processing/tree/main/MOFA/

### Dimension reduction analysis

The dimensions of the metabolomics and proteomics data sets were reduced to two/three dimensions using principal components analysis (PCA), t-distributed stochastic neighbour embedding (t-SNE) and Uniform Manifold Approximation and Projection (UMAP). Prior to performing PCA, the transformed and imputed intensity values were feature-wise scaled and centred before calculating PCs using prcomp (from the stats package, version 4.1.0). t-SNE was run using the Rtsne function and the following parameters: initial dimensions = 10, maximum number of iterations = 100, final dimensions = 3, perplexity = 3 (Rtsne package, version 0.15). UMAP was run using the umap function and the following parameters: minimum distance = 0.1, number of neighbours = 15, spread = 1 (umap package, version 0.2.7.0).

### Partial least square – discriminant analysis

To discriminate the samples based on the class vector Y, partial least square-discriminate analysis was performed. Y is here a vector of length n that indicates the class of each samples, i.e. a vector containing information on the *time*, *temperature*, *time*/*temperature* or the *individual* identifier. X is a n x p matrix containing the normalised+transformed+imputed intensities. To find the optimal number of components, plsda from the mixOmics package (version 6.15.45) was run with a maximum of 20 components (ncomp = 20), followed by evaluation of the performance of the fitted PLS using the perf function (validation = “Mfold”, folds = 3, nrepeat = 30). The overall classification error rate was taken as a measure to select the number of components and the number of components and distance method was selected by the maximum of the determined component number of the distances “centroids.dist”, “mahalanobis.dist” and “max.dist”. In a next step, the optimal number of variables was determined using a grid-based search ranging from 5 to 100 variables by the tune.splsda function (number of components, ncomp, and distance method, dist, as previously determined by the perf function, validation = “Mfold”, folds = 3, nrepeat = 30, measure = “BER”). The final model, using the optimal number of components based on t-tests on the error rate and the corresponding number of selected variables, was selected using the splsda function (scale = TRUE). All functions were taken from the mixOmics package (version 6.15.45). The code for the analysis can be found here: https://github.com/tnaake/SMARTCARE_preanalytical_processing/tree/main/MLM/metabolomics and https://github.com/tnaake/SMARTCARE_preanalytical_processing/tree/main/MLM/proteomics

### ‘Stability’ analysis using mixed linear models

For the mixed linear model, for plasma samples, the *time, temperature* and the interaction between *time* and *temperature* were included as fixed effects and individual as a random effect into the model. For serum samples, *time* was included as a fixed effect and individual as a random effect. When fitting the actual model, the lmer function from the lmerTest package (version 3.1-3) was used. If the mixed linear model was singular, an analysis of variance model was fitted with the same terms as for the mixed linear model except the random effects. The corresponding p-values to the fixed effects (*time*, *temperature*, and interaction between *time* and *temperature*) and the intercept were adjusted via FDR using the Benjamini-Hochberg method (α < 0.05). The code for the analysis can be found here: https://github.com/tnaake/SMARTCARE_preanalytical_processing/tree/main/MLM/metabolomics and https://github.com/tnaake/SMARTCARE_preanalytical_processing/tree/main/MLM/proteomics

### Pathway analysis of proteomic data

Pathway sets were obtained from the hallmark pathway set collection of Molecular Signature Database of the Broad Institute (MSigDB), with gene symbol identifiers. Pathway enrichment analysis was performed using the FGSEA R package (Version: fgsea_1.18.0), using the proteomic t-values (from **‘Stability’ analysis using mixed linear models** part) as input statistics. We set the number of permutations to 10.000 and only considered pathway sets with at least 10 protein members. All codes for this part is available at: https://github.com/saezlab/SMARTCARE_pilot_serum_prot_metab

### Proteomic and metabolomic signatures of plasma and serum samples

The pre-analytical quality signatures were made by selecting the top 20 p-values from the limma differential analysis output for serum and plasma samples stored for 8 h compared to 0 h on ice (4°C) or at RT. The contamination signatures were taken from Geyer *et al*. (2019). In order to estimate the normalised enrichment score, we used the weighted mean method from the decoupleR R package. The weights were either the t-values of the limma differential analysis for the storage quality signatures, the -1*t-values differences for the coagulation signature from Geyer *et al*. (2019) and 1 otherwise (for erythrocyte and platelet contamination, since no continuous weights were available). The signatures and method to estimate scores for plasma and serum samples are provided in the form of an open-source R package, that can be downloaded here: https://github.com/saezlab/plasmaContamination. The code for the analysis of the sample and computing the scores can be found here: https://github.com/saezlab/SMARTCARE_pilot_serum_prot_metab

## Supplementary Information

**Supplementary Figure 1:**
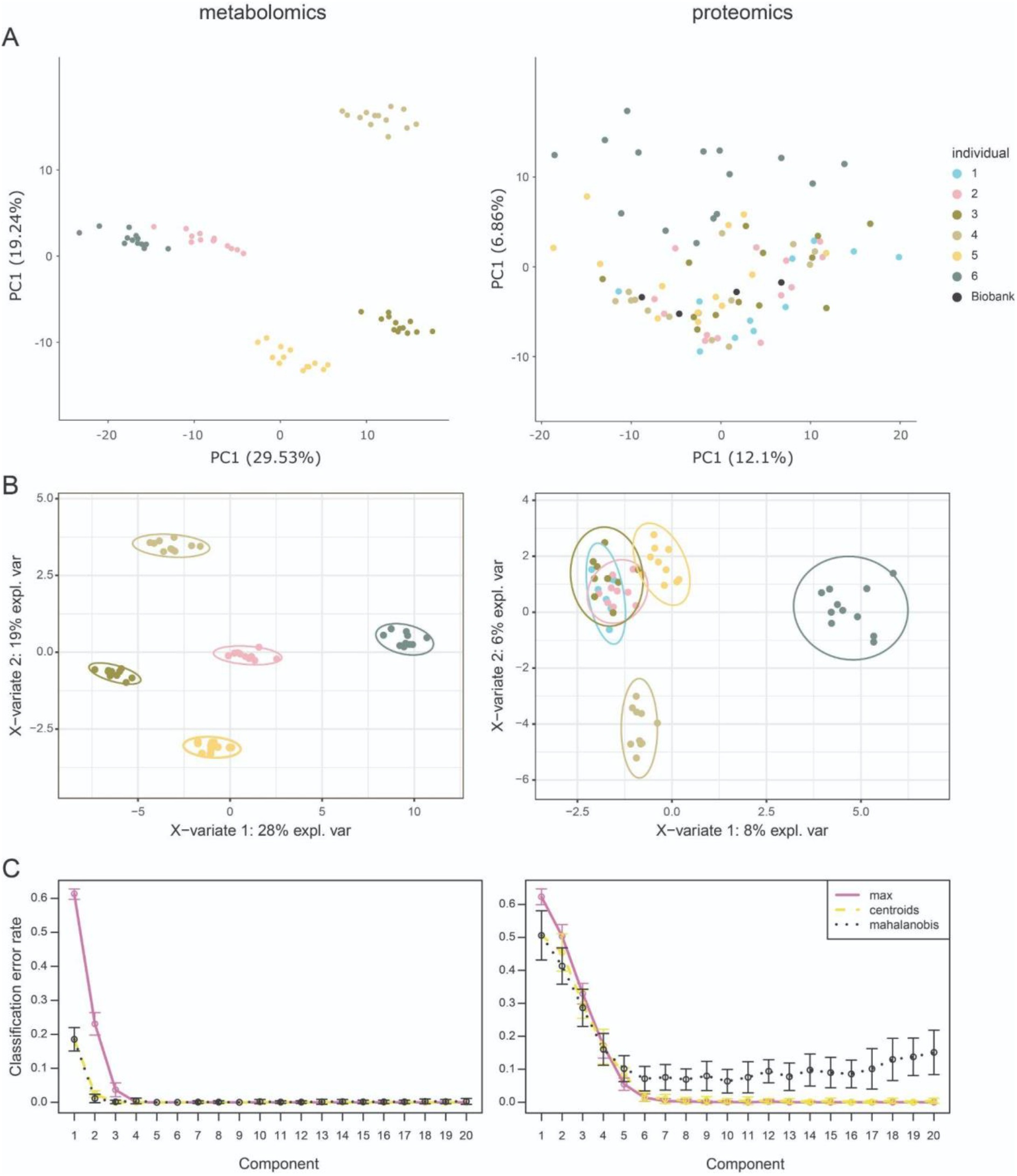
Individual-specific effects in the metabolomics and proteomics data set. A: Principal component analysis for metabolomics and proteomics data set identify individual-specific effects driving the variation within the individual data sets. The effect is more pronounced for the metabolomics data set. Biobank data points refer to long-term reference proteomics plasma samples. B: Sparse partial least squares – discriminant analysis classification using the set of features shown in Supplementary Table 1. The set of features leads to clear separation of individuals in the case of metabolomics, while for the proteomics data set a clear separation is not possible, indicating that it is more difficult to predict the individual based on proteins than on metabolites. C: Classification error rate of Partial least squares – discriminant analysis. Both analyses yield low classification error rates, indicating in general a separability according to the individuals.

**Supplementary Figure 2:**
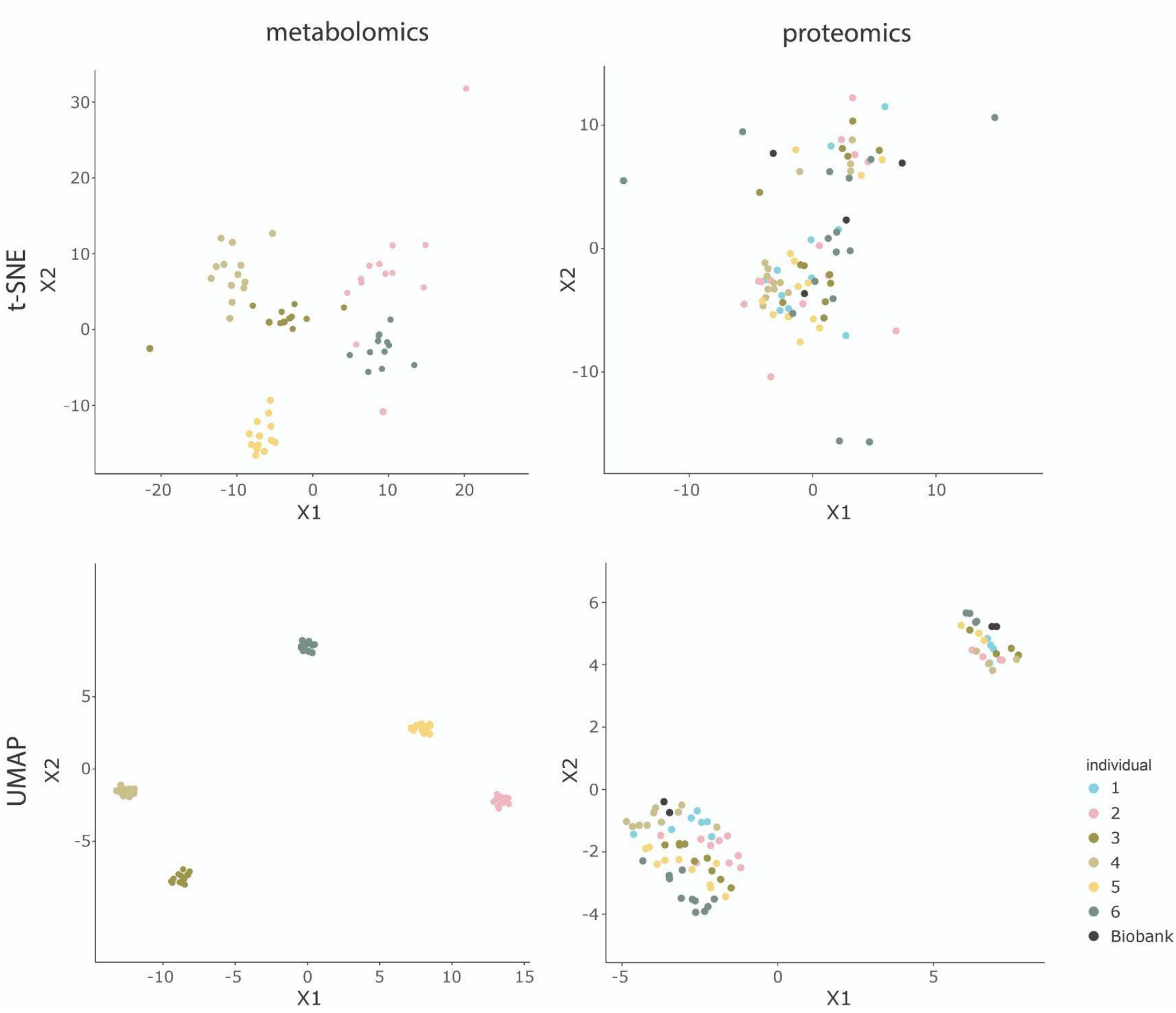
Dimension reduction analysis by t-SNE and UMAP of the metabolomics and proteomics data set. t-SNE and UMAP reveals for the metabolomics data set individual-specific effects, while for the proteomics data set t-SNE and UMAP separate the data according to plasma and serum. Biobank data points refer to long-term reference proteomics plasma samples.

**Supplementary Figure 3:**
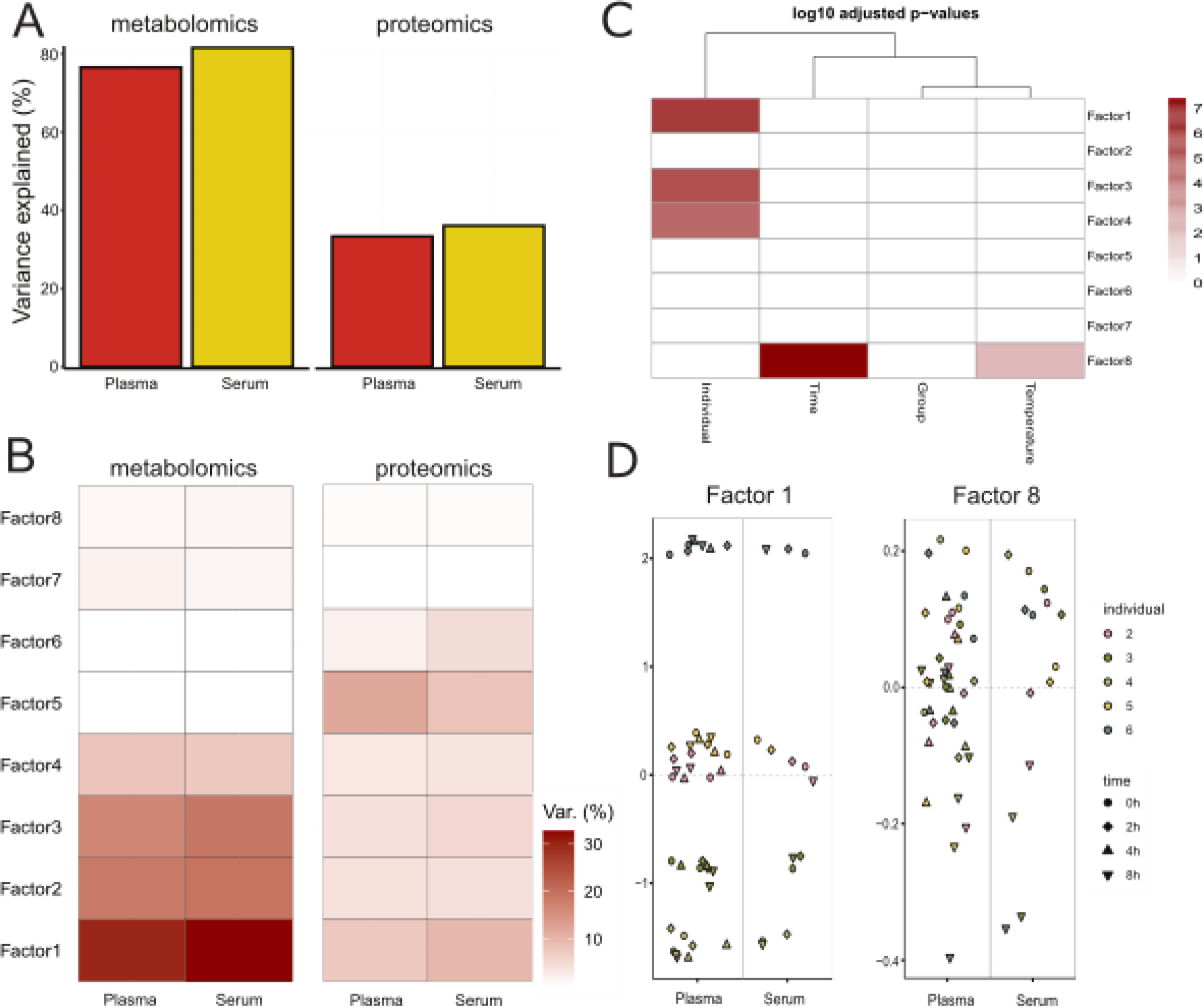
Multi-omics factor analysis (MOFA) of the metabolomics and proteomics data set. MOFA infers (hidden) factors that explain biological and technical sources of variability. The factors capture major sources of variation across the proteomics and metabolomics data sets (Argelaguet *et al*., 2018)). The MOFA model consisted of eight orthogonal axes of heterogeneity (factors 1 to 8) using the 40 and 14 joint samples for metabolomics and proteomics from plasma and serum. A: Variance explained for the metabolomics and proteomics data set for the plasma and serum groups. The fitted model explained 76.6 and 81.5% (R^2^_total_) of the variance in plasma and serum for the metabolomics data set, while for the proteomics data set, it explained 33.36% and 36.1% (R^2^_total_) of the variance in plasma and serum, respectively. B: Factor-wise explanation of variance. The metabolomics data set explains for most of the factors more variance than the proteomics data set. Factor 1 explains 29.9% and 7.5% (R^2^_total_) of the variance in plasma and 32.7% and 9.9% in serum for metabolomics and proteomics, respectively. Factor 8 explains 1.1% and 0.27% (R^2^_total_) of the variance in plasma and 1.32% and 0.47% (R^2^_total_) in serum for metabolomics and proteomics, respectively. C: Association test to check for the association of the eight factors with *individual*, *time*, *group* (plasma, serum), and *temperature*. Factors 1, 3, and 4 were associated with the *individual*, thereby explaining most of the variance within the data sets, and factor 8 was associated with the *time* variable. D: Visualisation of factors 1 and 8 in latent space. Factor 1 is separating the data set according to the *individual*, while factor 8 is separating the data set according to *time* and *temperature*. The variability along the x-axis includes random jitter.

**Supplementary Figure 4:**
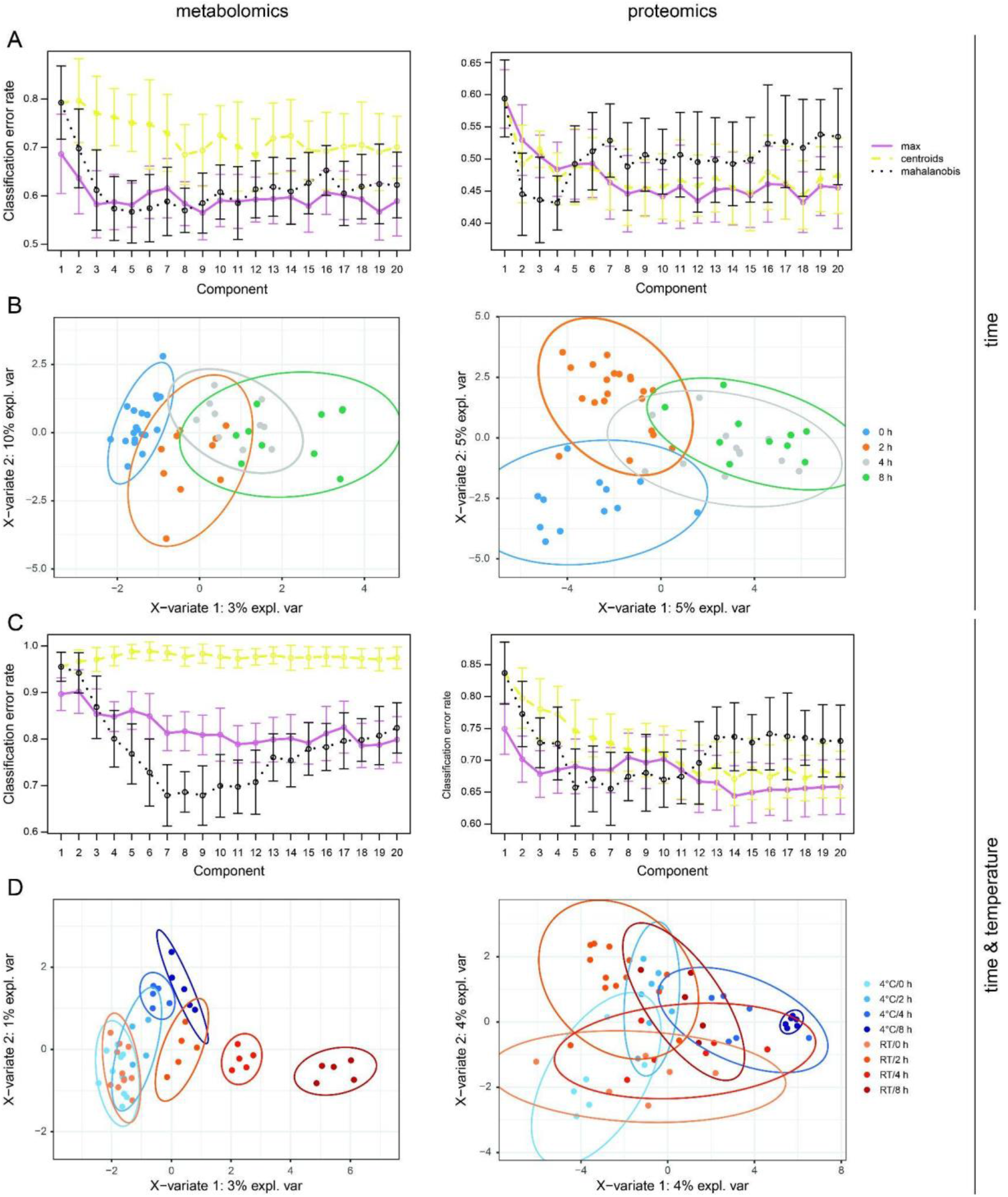
Partial least squares – discriminant analysis (PLS-DA) for the metabolomics and proteomics data sets. A: Classification error rates for PLS-DA using the class vector with *time* information. B: Sparse PLS-DA using the class vector with *time* information orders the samples along the time axis. C: Classification error rates for PLS-DA using the class vector with combined information on *time*/*temperature*. D: Sparse PLS-DA using the class vector with combined information on *time*/*temperature*. For the metabolomics data set, there are more distinct clusters for the samples subjected to RT incubation, while the clusters of the samples at 4°C are less distinct, indicating stronger effects on metabolite levels under RT. The sparse PLS-DA showed a more cluttered picture for the proteomics data set, indicating that the selected protein features are less suitable for classification of the combined *time/temperature* information.

**Supplementary Figure 5:**
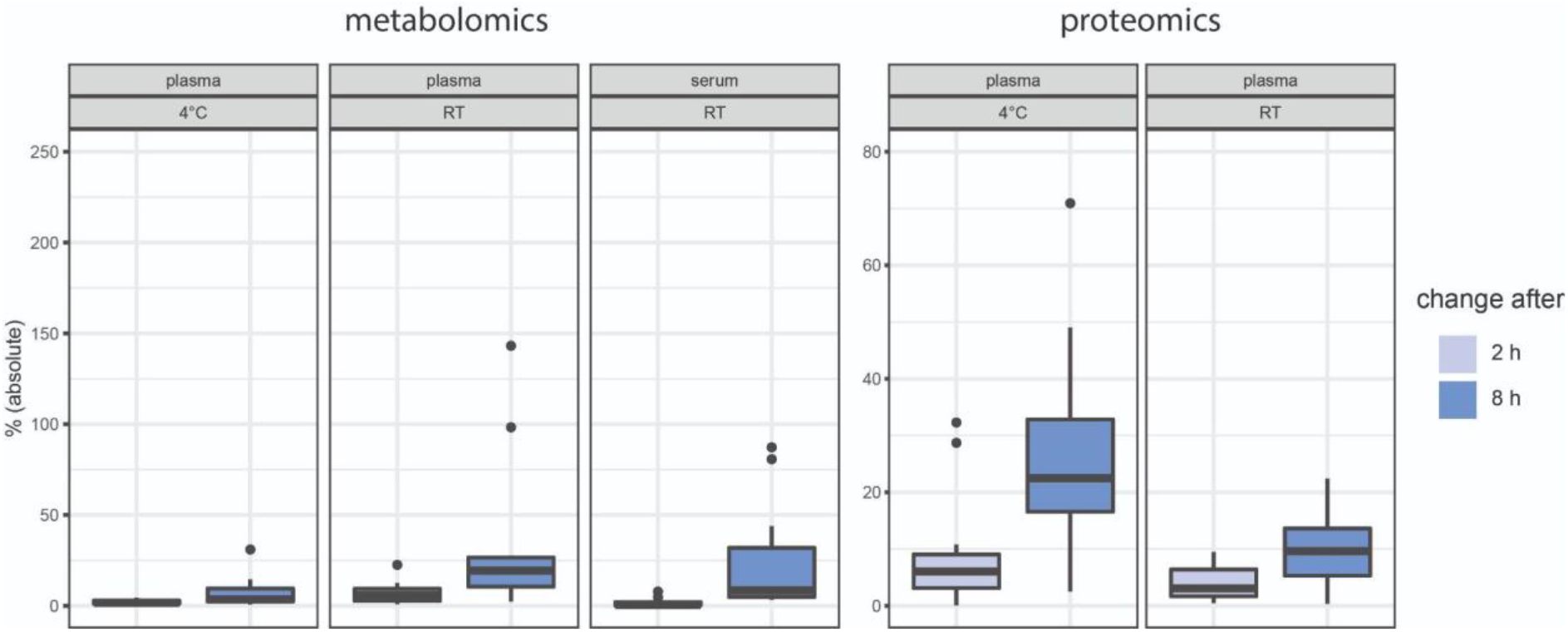
Absolute change of significantly changing metabolite and protein levels (in %) for the time points T2 h and T8 h compared to T0 h. The intensities at time point 0 h are set to 0 and the changes to the time point 2 h, 4 h, and 8 h are displayed as the mean changes of the individuals (in %). For plasma, the features are included that are significant to the pre-analytical factors *time*, *temperature* or the interaction *time*/*temperature* (α < 0.05, FDR correction). For serum, the features are included that are significant to the pre-analytical factor *time* (α < 0.05, FDR correction).

**Supplementary Figure 6:**
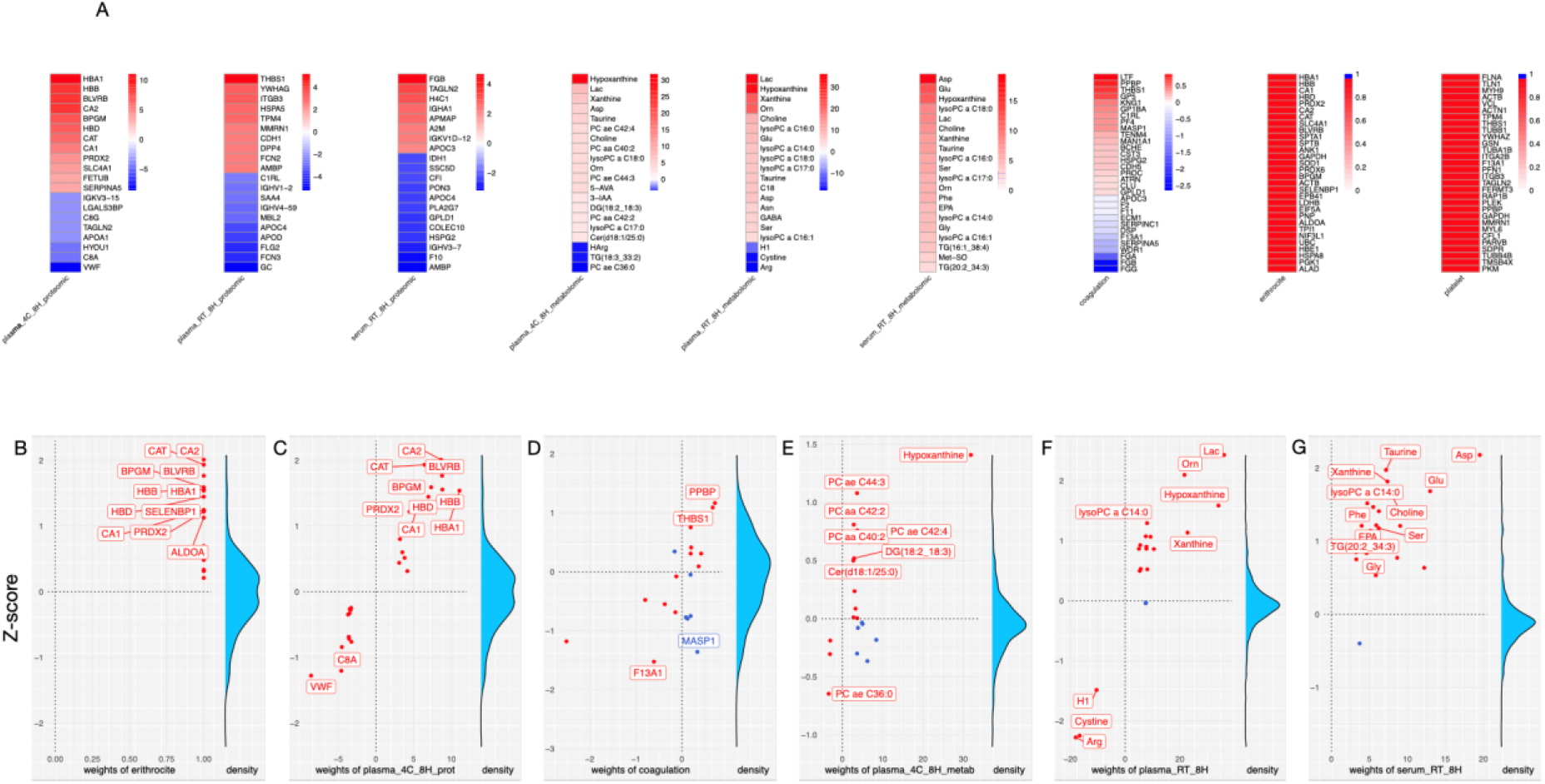
Details of signatures included in the plasmaContamination package. A: Proteins and metabolites composing the tested signatures. The colour represents the weight of each feature of the signature, when available. The signature score is computed as a weighted mean of the feature weight * feature measurement, normalised using an empirical distribution of score generated through feature shuffling. B, C, D, E, F, G: Scatter plot of feature weight against feature measurements for the corresponding samples/contrasts (a contrast being the result of a differential analysis between two conditions). B, C, E: relative to plasma samples kept at 4°C for 8 h. D, G: relative to serum samples kept at RT for 8 h. F: relative to plasma samples kept at RT for 8 h. Red and blue colours represent positive and negative contributions to the signature score, respectively. Red corresponds to up/down-regulated features with positive/negative weights, respectively, and blue corresponds to up/down-regulated features with negative/positive weights, respectively. The features that are the furthest away from the (0, 0) coordinate have the highest contribution to the score (contribution of a single feature being defined as weight * measurement).

**Supplementary Table 1:** Selected features of sparse partial least square – discriminant analysis using the *individual* as the class variable. The intensities of the features are ordered in descending order according to the absolute importance values.

| metabolite | Value of selected metabolite | protein | Value of selected protein |
| --- | --- | --- | --- |
| PC.ae.C36:5 | -0.268500768 | SAA2 (P0DJI9) | 0.58498936 |
| GDCA | -0.266503328 | C9 (P02748) | 0.51144060 |
| PC.ae.C34:3 | -0.256407863 | IGLV3-21 (P80748) | -0.37696364 |
| CE(14:0) | -0.229114859 | CRP (P02741) | 0.28526630 |
| PC.ae.C36:4 | -0.223906293 | PON1 (P27169) | -0.23718690 |
| CE(20:4) | -0.223493432 | APOC4 (P55056) | -0.18297455 |
| PC.ae.C38:5 | -0.218488828 | IGLV2-8 (P01709) | 0.17191216 |
| TG(20:2_34:1) | 0.218144999 | CFB (P00751) | 0.15198452 |
| PC.aa.C38:4 | -0.214187090 | F13A1 (P00488) | -0.14917795 |
| TG(20:1_34:2) | 0.207074762 | GSTO1 (P78417) | 0.04788159 |
| PC.ae.C38:4 | -0.204103703 | FCGBP (Q9Y6R7) | 0.04452931 |
| TG(16:0_38:2) | 0.191128128 | PRAP1 (Q96NZ9;Q96NZ9-2;Q96NZ9-3;Q96NZ9-4) | 0.04218922 |
| TG(20:1_34:1) | 0.173650357 | TTR (P02766) | -0.03921732 |
| CE(17:1) | -0.172349100 | C1S (P09871) | -0.01751029 |
| PC.ae.C32:1 | -0.170670334 | ALDOB (P05062) | -0.01389305 |
| PC.ae.C38:6 | -0.167923231 |  |  |
| PC.ae.C40:5 | -0.161312384 |  |  |
| HexCer(d18:2/24:0) | -0.159319689 |  |  |
| PC.aa.C36:4 | -0.158318121 |  |  |
| TDCA | -0.153837447 |  |  |
| HexCer(d18:1/24:0) | -0.143365272 |  |  |
| TG(16:0_38:3) | 0.132706459 |  |  |
| DCA | -0.119516370 |  |  |
| PC.aa.C40:5 | -0.110879600 |  |  |
| TG(18:1_36:1) | 0.109261217 |  |  |
| GUDCA | 0.101513121 |  |  |
| lysoPC.a.C20:4 | -0.094747816 |  |  |
| PC.aa.C38:0 | -0.086969074 |  |  |
| PC.ae.C34:2 | -0.086936894 |  |  |
| TG(18:0_36:3) | 0.081121569 |  |  |
| PC.ae.C40:6 | -0.080362096 |  |  |
| CE(22:5) | -0.079775441 |  |  |
| TG(18:0_36:2) | 0.077673974 |  |  |
| FA(18:2) | 0.074525414 |  |  |
| CE(18:0) | -0.069044645 |  |  |
| TG(18:1_34:2) | 0.046089288 |  |  |
| HexCer(d18:1/23:0) | -0.043501804 |  |  |
| SM.C18:1 | -0.040112031 |  |  |
| PC.aa.C38:5 | -0.034541815 |  |  |
| HipAcid | -0.033557136 |  |  |
| CE(15:0) | -0.027731165 |  |  |
| Kynurenine | 0.027723492 |  |  |
| CE(14:1) | -0.026967113 |  |  |
| TrpBetaine | 0.026213738 |  |  |
| PC.aa.C40:4 | -0.024257807 |  |  |
| lysoPC.a.C28:1 | -0.022013954 |  |  |
| PC.ae.C32:2 | -0.012042735 |  |  |
| CE(18:1) | -0.003493789 |  |  |
| CE(16:1) | -0.001702684 |  |  |
| PC.ae.C34:1 | -0.000103610 |  |  |

**Supplementary Table 2:** Selected features of sparse partial least square – discriminant analysis using the *time* as the class vector. The features are ordered in descending order according to the absolute importance values.

| Metabolite | Value of selected metabolite | Protein | Value of selected protein |
| --- | --- | --- | --- |
| Hypoxanthine | 0.86500382 | HBA1 (P69905) | 0.3051776936 |
| Xanthine | 0.35585116 | ECM1 (Q16610;Q16610-4) | -0.2790432656 |
| Lactate | 0.33061249 | BPGM (P07738) | 0.2728424861 |
| Ornithine | 0.12100160 | APOA1 (P02647) | -0.2689895678 |
| Asparagine | 0.03453019 | CAT (P04040) | 0.2574125644 |
|  |  | CA2 (P00918) | 0.2506049009 |
|  |  | HYOU1 (Q9Y4L1) | -0.2468780420 |
|  |  | SLC4A1 (P02730) | 0.2259034325 |
|  |  | FERMT3 (Q86UX7;Q86UX7-2) | 0.2201556911 |
|  |  | HBB (P68871) | 0.2103457860 |
|  |  | BASP1 (P80723;P80723-2) | -0.2062149021 |
|  |  | HBD (P02042) | 0.1863074451 |
|  |  | BLVRB (P30043) | 0.1823127035 |
|  |  | CA1 (P00915) | 0.1754204972 |
|  |  | YWHAZ (P63104) | 0.1705790612 |
|  |  | MPO (P05164;P05164-2;P05164-3) | 0.1635279760 |
|  |  | GANAB (Q14697;Q14697-2) | 0.1397937894 |
|  |  | GC (P02774) | -0.1394798788 |
|  |  | F2 (P00734) | -0.0955141175 |
|  |  | C8G (P07360) | -0.0934011313 |
|  |  | CFL1 (P23528) | 0.0886430466 |
|  |  | PRSS3 (P35030;P35030-2;P35030-3;P35030-4) | 0.0864709428 |
|  |  | PROZ (P22891;P22891-2) | 0.0792679857 |
|  |  | FN1 (P02751;P02751-3) | -0.0788933328 |
|  |  | HPX (P02790) | -0.0709280449 |
|  |  | VASN (Q6EMK4) | -0.0665273047 |
|  |  | ATRN (O75882-2;O75882-3) | 0.0650318704 |
|  |  | PSMA4 (P25789) | -0.0628043092 |
|  |  | KRT8 (P05787) | -0.0606475289 |
|  |  | MDH1 (P40925;P40925-2) | 0.0599833383 |
|  |  | IGKV3-20 (P04206) | -0.0587456769 |
|  |  | EEF1A1 (P68104;Q5VTE0) | -0.0584839501 |
|  |  | PRDX2 (P32119) | 0.0578632531 |
|  |  | ITIH2 (P19823) | 0.0550239670 |
|  |  | PTPRG<br>(P23470;P23470-2) | 0.0438356128 |
|  |  | EFEMP1<br>(Q12805;Q12805-2;Q12805-3;Q12805-4) | 0.0429530829 |
|  |  | ITIH3 (Q06033;Q06033-2) | 0.0423345831 |
|  |  | ITGB3 (P05106) | 0.0414866691 |
|  |  | HNRNPK<br>(P61978;P61978-2;P61978-3) | -0.0407078585 |
|  |  | ALDOA (P04075) | 0.0406151070 |
|  |  | GP1BB<br>(P13224;P13224-2) | 0.0405218107 |
|  |  | TPM3 (P06753-2) | 0.0397427929 |
|  |  | IGLC2 (P0CG05) | -0.0388841132 |
|  |  | IGHG1 (P01857) | 0.0387823216 |
|  |  | TFRC (P02786) | 0.0383223128 |
|  |  | VWF (P04275) | -0.0332107407 |
|  |  | IGHV3-7 (P80419) | -0.0331100294 |
|  |  | MRC1 (P22897) | 0.0325502649 |
|  |  | HSPA8<br>(P11142;P11142-2) | 0.0324084800 |
|  |  | PRDX1 (Q06830) | -0.0292653697 |
|  |  | PGK1 (P00558) | 0.0266566339 |
|  |  | F12 (P00748) | 0.0246458975 |
|  |  | FGL1 (Q08830) | 0.0237648829 |
|  |  | CFI (P05156) | 0.0230229098 |
|  |  | CLU<br>(P10909;P10909-2;P10909-4;P10909-5) | 0.0223664138 |
|  |  | PRDX6 (P30041) | 0.0182668762 |
|  |  | IGLV6-57 (P06318) | 0.0160594551 |
|  |  | FLT4<br>(P35916;P35916-1;P35916-3) | -0.0159933947 |
|  |  | TLN1 (Q9Y490) | 0.0143133844 |
|  |  | APOA4 (P06727) | -0.0138128179 |
|  |  | MASP1<br>(P48740-2;P48740-4) | 0.0130026995 |
|  |  | NME2 (P22392;P22392-2) | 0.0122881335 |
|  |  | ITGA2B (P08514;P08514-2) | 0.0109002438 |
|  |  | AHSG (P02765) | -0.0106803269 |
|  |  | ANGPTL3 (Q9Y5C1) | -0.0102464019 |
|  |  | AFM (P43652) | -0.0091812995 |
|  |  | DPP4 (P27487) | 0.0081835342 |
|  |  | IGKV1D-33 (P01593) | -0.0055046478 |
|  |  | KNG1 (P01042-2) | 0.0042764859 |
|  |  | DSG2 (Q14126) | 0.0037493018 |
|  |  | C8A (P07357) | -0.0035858940 |
|  |  | NFX1 (Q12986;Q12986-2) | -0.0031747150 |
|  |  | DSC1 (Q08554;Q08554-2) | -0.0030473481 |
|  |  | HLA-A<br>(P01891;P01892;P04439;<br>P05534;P10314;P10316;<br>P13746;P13746-2;<br>P16188;P16189;P16190;<br>P18462;P30443;P30447;<br>P30450;P30453;P30455;<br>P30456;P30457;P30459;<br>P30508;P30512;Q29960;<br>Q29960-2;Q95604) | 0.0013763727 |
|  |  | SAA4 (P35542) | -0.0002944848 |

**Supplementary Table 3:** Selected features for sparse partial least square – discriminant analysis using *temperature* as the class vector. The features are ordered in descending order according to the absolute importance values.

| Metabolite | Value of selected metabolite | Protein | Value of selected protein |
| --- | --- | --- | --- |
| Lactate | -0.60085947 | THBS1 (P07996) | 0.6521617731 |
| Ornithine | -0.54910269 | PF4 (P02776;P10720) | 0.6318854387 |
| Xanthine | -0.36905947 | VWF (P04275) | 0.3135767399 |
| Arginine | 0.28784931 | AMBP (P02760) | 0.1908734707 |
| Cystine | 0.27148684 | PPBP (P02775) | 0.1750974544 |
| Aconitic acid | -0.20830526 | FN1 (P02751;P02751-3) | 0.0964168595 |
| Hexosyl ceramide (d18:2/16:0) | 0.03599758 | TKT (P29401) | -0.0231791401 |
|  |  | KRT6B (P04259) | -0.0122508252 |
|  |  | GP1BB (P13224;P13224-2) | -0.0002753722 |

**Supplementary Table 4:**
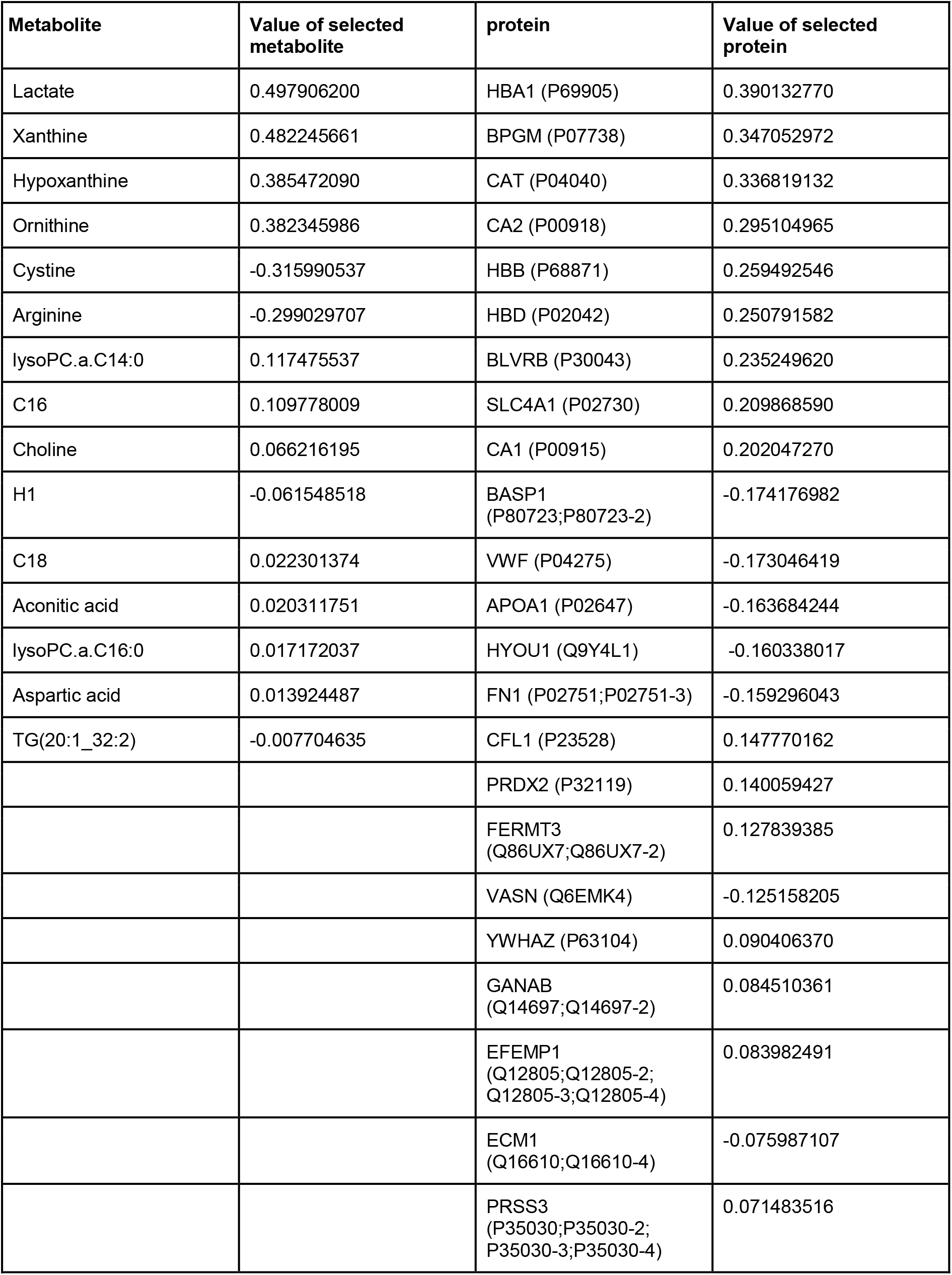
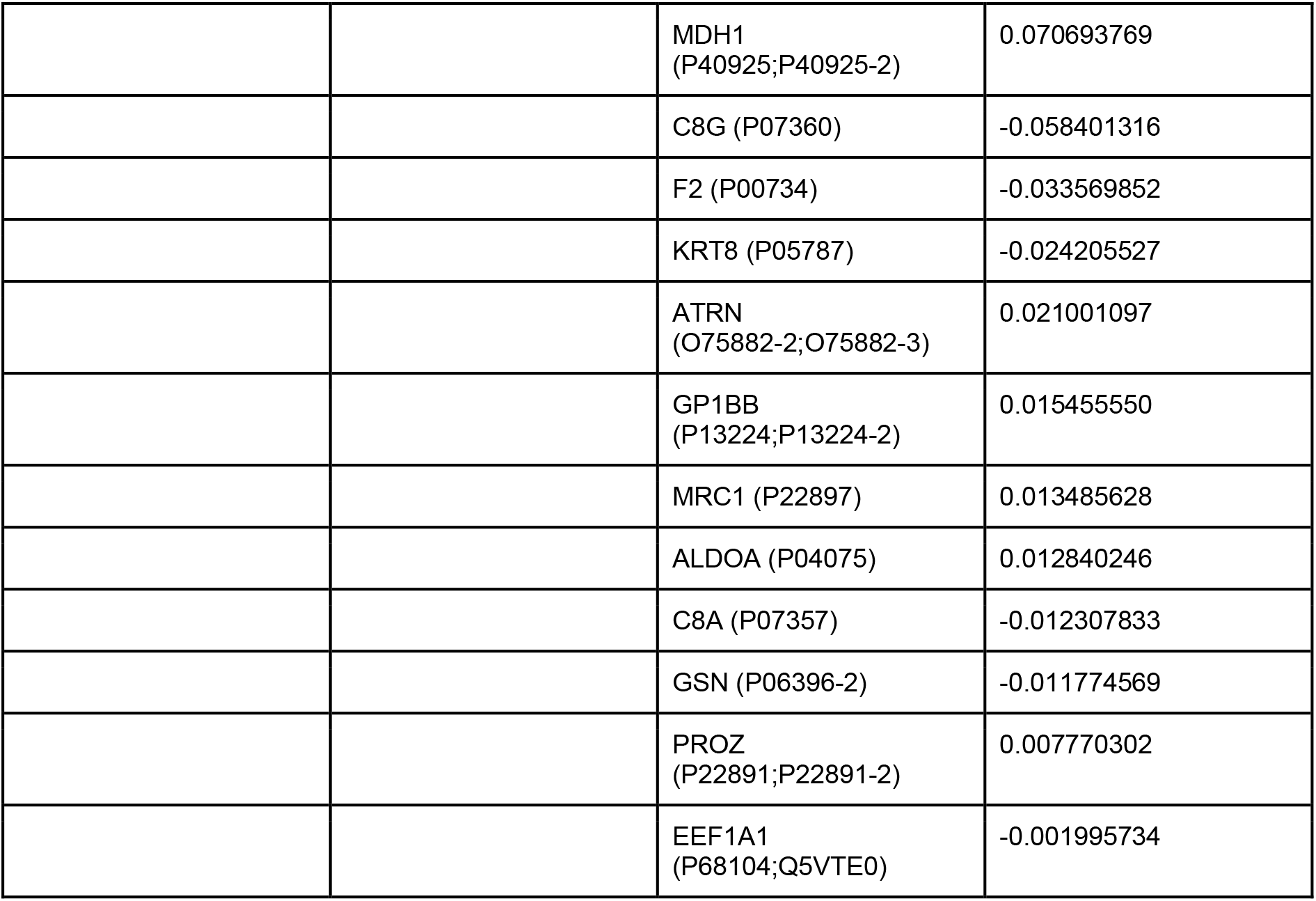
Selected features for spare partial least square – discriminant analysis using the combined factor *time/temperature* as the class vector. The features are ordered in descending order according to the absolute importance values.

| Metabolite | Value of selected metabolite | protein | Value of selected protein |
| --- | --- | --- | --- |
| Lactate | 0.497906200 | HBA1 (P69905) | 0.390132770 |
| Xanthine | 0.482245661 | BPGM (P07738) | 0.347052972 |
| Hypoxanthine | 0.385472090 | CAT (P04040) | 0.336819132 |
| Ornithine | 0.382345986 | CA2 (P00918) | 0.295104965 |
| Cystine | -0.315990537 | HBB (P68871) | 0.259492546 |
| Arginine | -0.299029707 | HBD (P02042) | 0.250791582 |
| lysoPC.a.C14:0 | 0.117475537 | BLVRB (P30043) | 0.235249620 |
| C16 | 0.109778009 | SLC4A1 (P02730) | 0.209868590 |
| Choline | 0.066216195 | CA1 (P00915) | 0.202047270 |
| H1 | -0.061548518 | BASP1 (P80723;P80723-2) | -0.174176982 |
| C18 | 0.022301374 | VWF (P04275) | -0.173046419 |
| Aconitic acid | 0.020311751 | APOA1 (P02647) | -0.163684244 |
| lysoPC.a.C16:0 | 0.017172037 | HYOU1 (Q9Y4L1) | -0.160338017 |
| Aspartic acid | 0.013924487 | FN1 (P02751;P02751-3) | -0.159296043 |
| TG(20:1_32:2) | -0.007704635 | CFL1 (P23528) | 0.147770162 |
|  |  | PRDX2 (P32119) | 0.140059427 |
|  |  | FERMT3 (Q86UX7;Q86UX7-2) | 0.127839385 |
|  |  | VASN (Q6EMK4) | -0.125158205 |
|  |  | YWHAZ (P63104) | 0.090406370 |
|  |  | GANAB (Q14697;Q14697-2) | 0.084510361 |
|  |  | EFEMP1 (Q12805;Q12805-2;Q12805-3;Q12805-4) | 0.083982491 |
|  |  | ECM1 (Q16610;Q16610-4) | -0.075987107 |
|  |  | PRSS3 (P35030;P35030-2;P35030-3;P35030-4) | 0.071483516 |
|  |  | MDH1<br>(P40925;P40925-2) | 0.070693769 |
|  |  | C8G (P07360) | -0.058401316 |
|  |  | F2 (P00734) | -0.033569852 |
|  |  | KRT8 (P05787) | -0.024205527 |
|  |  | ATRN<br>(O75882-2;O75882-3) | 0.021001097 |
|  |  | GP1BB<br>(P13224;P13224-2) | 0.015455550 |
|  |  | MRC1 (P22897) | 0.013485628 |
|  |  | ALDOA (P04075) | 0.012840246 |
|  |  | C8A (P07357) | -0.012307833 |
|  |  | GSN (P06396-2) | -0.011774569 |
|  |  | PROZ<br>(P22891;P22891-2) | 0.007770302 |
|  |  | EEF1A1<br>(P68104;Q5VTE0) | -0.001995734 |

## Author contributions

F.C. and H.M.G. designed the experiment. F.C., H.M.G. and N.K.-R. performed the experiment. H.M.G., G.K. and T.M. measured the metabolites and proteins. T.N. and A.D. analysed and interpreted the data set. A.D., H.M.G., T.M., and T.N. wrote the manuscript. E.J. and C.H suggested methods and experiments and provided feedback on the manuscript. B.H., U.K., C.M.-T., S.D., J.S.-R., W.H., R.H., G.P., and J.K. provided feedback to the manuscript.

## Conflict of Interest

J.S.-R. reports funding from GSK and Sanofi and fees from Travere Therapeutics and Astex Therapeutics.

## Acknowledgements

This work was supported by the German Ministry of Education and Research (BMBF), as part of the National Research Node “Mass spectrometry in Systems Medicine” (MSCoreSys), under the funding code 161L0212. Several figures were created with BioRender.com.

